# Interconnected feedback loops among ESRP1, HAS2, and CD44 regulate epithelial-mesenchymal plasticity in cancer

**DOI:** 10.1101/260349

**Authors:** Mohit Kumar Jolly, Bogdan-Tiberius Preca, Satyendra C Tripathi, Dongya jia, Samir M Hanash, Thomas Brabletz, Marc P Stemmler, Jochen Maurer, Herbert Levine

## Abstract

Aberrant activation of epithelial-mesenchymal transition (EMT) in carcinoma cells contributes to increased migration and invasion, metastasis, drug resistance, and tumor-initiating capacity. EMT is not always a binary process, rather cells may exhibit a hybrid epithelial/mesenchymal (E/M) phenotype. ZEB1 - a key transcription factor driving EMT - can both induce and maintain a mesenchymal phenotype. Recent studies have identified two novel autocrine feedback loops utilizing ESRP1, HAS2, and CD44 that maintain high levels of ZEB1. However, how the crosstalk between these feedback loops alters the dynamics of epithelial-hybrid-mesenchymal transition remains elusive. Here, using an integrated theoretical-experimental framework, we identify that these feedback loops can enable cells to stably maintain a hybrid E/M phenotype. Moreover, computational analysis identifies the regulation of ESRP1 as a crucial node, a prediction that is validated by two complementary experiments showing that (a) overexpression of ESRP1 reverts EMT in MCF10A cells treated with TGFβ for 21 days, and (b) knockdown of ESRP1 in stable hybrid E/M H1975 cells drives EMT. Finally, in multiple breast cancer datasets, high levels of ESRP1, ESRP1/HAS2, and ESRP1/ZEB1 correlates with poor prognosis, supporting the relevance of ZEB1/ESRP1 and ZEB1/HAS2 axes in tumor progression. Together, our results unravel how these interconnected feedback loops act in concert to regulate ZEB1 levels and to drive the dynamics of epithelial-hybrid-mesenchymal transition.

## Introduction

Epithelial-Mesenchymal Transition (EMT) and its reverse Mesenchymal-Epithelial Transition (MET) are embryonic programs that get aberrantly activated during tumor progression [1],. and regulate metastasis [2]. EMT is characterized by affecting one or more of these traits - decreased cell-cell adhesion rearranged cytoskeleton disrupted apico-basal polarity and increased migration and invasion [3]. Furthermore, EMT contributes to drug resistance [4], evasion of the immune system [5], and tumor-initiating properties [67]. EMT can be induced by multiple micro-environmental conditions such as matrix stiffness [8,9] and hypoxia [10] that can activate one or more of EMT-driving transcription factors such as ZEB (ZEB1/2), Snail (SNAI1/2), and Twist [11]. ZEB1 - a key driver of EMT - can confer tumor-initiating (stemness) and chemoresistance properties to cancer cells [12, 13], and its expression correlates with poor prognosis [14]. ZEB1 mainly acts as a transcriptional repressor for genes involved in cell-cell adhesion and cell polarity [11], and for microRNAs that promote an epithelial phenotype such as the miR-200 family [15,16],. The miR-200 family also represses ZEB1 translation thereby forming a mutually inhibitory feedback loop [17, 18], referred to as the ‘motor of cellular plasticity’ [19]. Higher levels of ZEB1 is key to maintaining a mesenchymal phenotype [20].

Recent computational models have suggested that the miR-200/ZEB1 feedback loop can not only allow cells to attain an epithelial (high miR-200, low ZEB) and a mesenchymal (low miR-200, high ZEB) phenotype [21,22], but also a hybrid epithelial/mesenchymal (E/M) state - (medium miR-200, medium ZEB) [23–25]. Indeed, active nuclear ZEB1 was observed in H1975 lung cancer cells [20] that maintain a stable hybrid E/M state at a single-cell level over many passages [27]. These earlier models presumed a self-activation of ZEB1 [23–25] based on ZEB1-mediated stabilization of SMAD complexes [23] (Figure 1A). More recently exact molecular mechanisms mediating a ZEB1 self-activation were elucidated [28,29].

**Figure 1.**
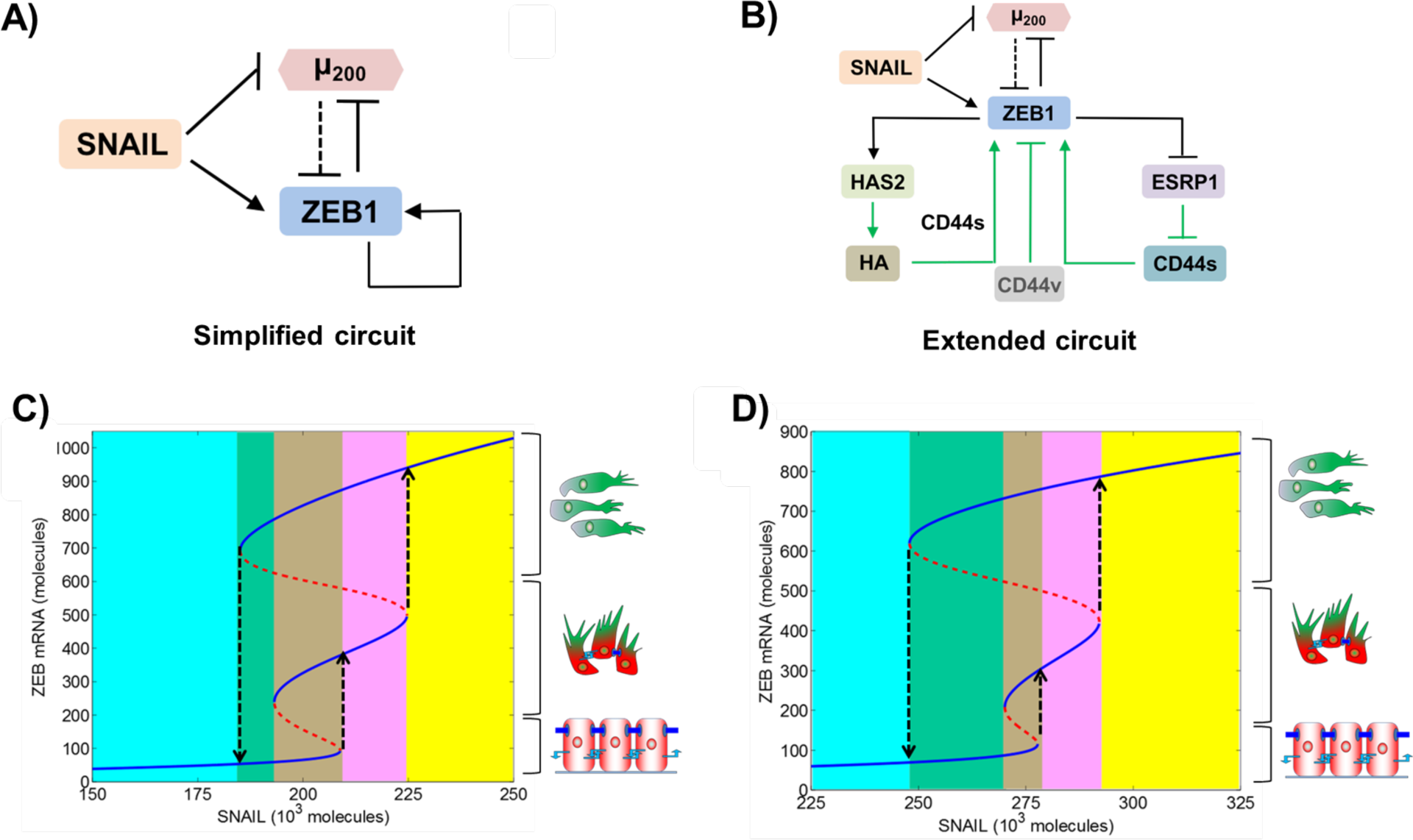
Dynamics of EMT regulatory circuit. **A)** Simplified EMT regulatory circuit - mutually inhibitory feedback loop between ZEB1 and miR-200, driven by SNAIL. In this earlier modeling attempt, selfactivation of ZEB1 is considered to be direct, to incorporate the stabilization of SMADs by ZEB1. ***B)*** Extended EMT regulatory circuit with explicit mechanistic details of ZEB1 self-activation, as identified in recent studies. Solid black lines show transcriptional regulation, dotted black lines show microRNA-mediated regulation, and green arrows indicate regulation whose modes are either non-transcriptional or yet have to be identified. ***C)*** Bifurcation diagram of the miR-200/ZEB1 (simplified) circuit shown in A), driven by SNAIL. Solid blue curves show stable states (cellular phenotypes), red dotted curves show unstable states. Low levels of ZEB1 denote an epithelial (E) phenotype, high levels of ZEB1 denote a mesenchymal (M) phenotype, and intermediate levels of ZEB1 denote a hybrid E/M phenotype, as shown by cartoons drawn alongside. Different colored regions show different phases - cyan region depicts {E} phase, green region depicts {E, M} phase, brown region depicts {E, E/M, M} phase, pink region depicts {E/M, M} phase, and yellow region depicts {M} phase. ***D)*** Bifurcation diagram for the extended, i.e. miR-200/ZEB1/ESRP1/HAS2/CD44 circuit as shown in B).

Particulary ZEB1 can self-activate through two interconnected feedback loops. First ZEB1 represses epithelial splicing regulatory protein 1 (ESRP1) [29] that controls alternative splicing of the cell surface receptor CD44 which is differentially regulated during EMT [30]. Decreased ESRP1 leads to enhanced levels of CD44s (CD44 standard; devoid of all variable exons) isoform, and depleted levels of variable-exon containing CD44v (CD44 variant) isoforms. Overexpression of CD44s, in turn activates ZEB1 whereas increase in CD44v isoforms inhibits ZEB1 [29] (Figure 1B). Second, ZEB1 activates hyaluronic acid synthase 2 (HAS2) that synthesizes hyaluronic acid (HA) - a key proteoglycan in the extracellular matrix that in turn elevates the levels of ZEB1 [20] (Figure 1B). These two feedback loops are connected due to interactions of HA with its receptor CD44. CD44s was reported to be essential for the effect of HA on ZEB1 suggesting that HA binding to its receptor CD44s affects EMT [20]. CD44s-HA interactions have been proposed to protect cells undergoing EMT against anoikis [31], a trait commonly associated with EMT [32], thereby reinforcing the role of CD44s-HA interactions in this process. CD44s overexpression together with excess of HA leads to even higher levels of ZEB1 as compared to CD44s overexpression alone [20]. Hence, CD44s may be able to activate ZEB1 through both HA-dependent and HA-independent pathways. However how these complex interconnections among these feedback loops regulating ZEB1 drive the emergent dynamics of epithelial-hybrid-mesenchymal transitions remains elusive.

Here, using an integrated theoretical-experimental approach we elucidated the dynamics of the interconnected feedback loops among ZEB1, ESRP1, HAS2, and CD44 in mediating epithelial-hybrid-mesenchymal transitions. First we constructed a mathematical model denoting the known interactions among these players. The model predicted that these feedback loops can enable for the existence of a stable hybrid E/M phenotype, besides epithelial and mesenchymal phenotypes, Sensitivity analyses on model parameters identify ESRP1 as a key mediator of EMT and MET. Consistenty overexpression of ESRP1 in MCF10A cells treated with TGFβ reversed EMT while knockdown of ESRP1 in stable hybrid E/M cells - H1975 - drove EMT. Finally, higher levels of ESRP1, ESRP1/HAS2, and ESRP1/ZEB1 correlates with poor patient survival, indicating the functional relevance of these feedback loops during tumor progression.

## Results

### Mathematical modeling suggests how the miR-200/ZEB1/ESRP1/HAS2/CD44 circuit drives the dynamics of epithelial-hybrid-mesenchymal transition

As a first step towards elucidating the dynamics of epithelial-hybrid-mesenchymal transition as driven by ZEB1/ESRP1/HAS2/CD44 feedback loops, we construct a mathematical model representing the experimentally known regulatory interactions among these players (SI section 1). Next we plot the levels of ZEB1 as a function of an EMT-inducing signal (here, represented by SNAIL), as predicted by the model. We also compared the dynamics of these transitions as mediated by the miR-200/ZEB1/ESRP1/HAS2/CD44 circuit with the control case, i.e. when ZEB1 self-activation is not included through these detailed pathways (miR-200/ZEB1 circuity but instead as in earlier modeling attempts [23–25].

The response of the different circuits - miR-200/ZEB1 (hereafter referred to as the ‘simplified circuit’) and miR-200/ZEB1/ESRP1/HAS2/CD44 (hereafter referred to as the ‘extended circuit’) - to varying levels of SNAIL (mimicking an induction of EMT) is presented as bifurcation diagrams of ZEB1 mRNA levels (Figure 1C D). Lower levels of ZEB1 mRNA (<150 molecules) denote an epithelial (E) phenotype, higher values (> 600 molecules) represent a mesenchymal (M) phenotype, and intermediate values (~200-500 molecules) denote a hybrid E/M phenotype (solid blue lines in Figure 1C D).

The dynamics of these two circuits look remarkably similar. For low SNAIL levels, cells maintain an E phenotype. Increase in SNAIL levels drive cells toward a hybrid E/M phenotype, and a further increase induces a complete EMT causing cells to attain an M state (black dotted upward arrows in Figure 1C D). Moreover for certain SNAIL levels, cells can exhibit more than one phenotype and thus can spontaneously interconvert among one another for instance, among E and M states in the {E, M} phase (green shaded region in Figure 1C D) among E, hybrid E/M and M states in the {E, E/M, M} phase (brown shaded region in Figure 1C D) and among hybrid E/M and M states in the {E/M, M} phase (pink shaded region in Figure 1 C D). These results suggest that ZEB1 self-activation mediated by ESRP1/CD44 and HAS2/HA/CD44 axes should be considered integral to the dynamics of epithelial-hybrid-mesenchymal transition.

Moreover in pan-cancer cell line cohorts NCI-60 and CCLE (Cancer Cell Line Encyclopedia) ZEB1 positively correlates with HAS2, and negatively correlates with ESRP1 [26,28,29]. This correlation is also observed in lung, breast and pancreatic tumors [28,29], suggesting that these interconnected feedback loops may regulate EMT across cancer types. This idea is further strengthened by a positive correlation between HAS2 and CD44s in breast cancer [33].

### Sensitivity analysis identifies ESRP1 as a key mediator of epithelial-hybrid-mesenchymal transitions

To evaluate the robustness of the model predictions mentioned above, we performed a sensitivity analysis to parameter perturbation. Our sensitivity analysis for the simplified circuit (i.e. miR-200/ZEB1 circuit without HAS2, CD44, and ESRP1) identified ZEB1 self-activation as a crucial link. Modulating the strength of self-activation significantly affected the range of SNAIL levels for which cells could acquire a stable hybrid E/M phenotype [25], consistent with observations for such feedback loops enabling three states [34,35].

To further assess which link(s) in the extended circuit (miR-200/ZEB1/ESRP1/HAS2/CD44) impact the stability of hybrid E/M phenotype the most, we varied every parameter in the model for this circuit by +/- 10%, one at a time, and calculated its effect on the range of SNAIL levels for which cells can acquire a hybrid E/M phenotype (highlighted by black rectangle in Figure 2A). While the model is largely robust to parameter variation, in a few cases, we noticed a relatively larger change in the region corresponding to a stable hybrid E/M phenotype (highlighted by arrows in Figure 2B). Particularly, when the strength of inhibition of ESRP1 by ZEB1 is increased, the range of SNAIL levels for the existence of hybrid E/M decreased (Table S1). Conversely, this range increased when the inhibition of ESRP1 by ZEB1 was decreased, or when the biosynthesis rates of ZEB1 were altered (Section S1). These results are in conceptual agreement with the sensitivity analysis for the simplified circuit [25]. Thus, the strength of ZEB1-ESRP1 interaction is likely to play a key role in controlling the stability of a hybrid E/M phenotype, and the dynamics of EMT and MET.

**Figure 2.**
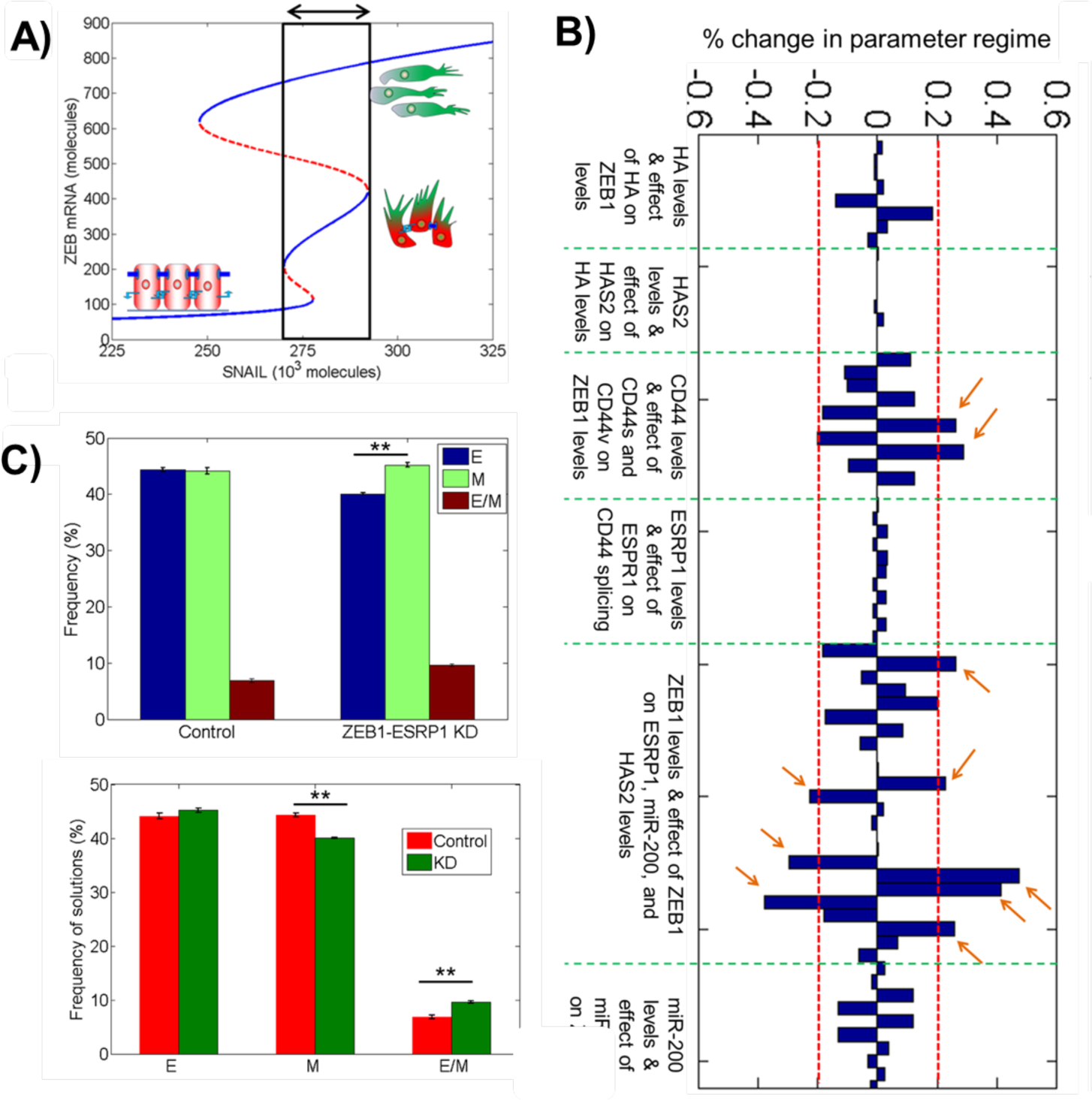
Sensitivity analysis identifies ZEB1-ESRP1 link as a key regulator for EMT. **A)** Bifurcation diagram shown in Figure 1D, highlighting the region for the existence of a hybrid E/M phenotype, either alone or in combination with other phenotypes. ***B)*** Sensitivity analysis for miR-200/ZEB1/ESRP1/HAS2/ CD44 circuit (Figure 1B), where the change in the width of the area highlighted in A) is plotted for +/- 10% of every parameter, one at a time. Parameters are grouped based on their role in the circuit; green dotted lines denote different groups as mentioned below each group. Red dotted lines marks over +/- 20% change in the width of the area highlighted in A). The model predictions are most sensitive to parameters that modulate the strength of ZEB1-ESRP1 link (marked by arrows) ***C)*** Solutions from an ensemble of 10,000*3 (n=3) mathematical models, each with a distinct set of parameters, for two conditions - with the ZEB1-ESRP1 link intact (control), and with ZEB1-ESRP1 node deleted (ZEB1-ESRP1 KD). ** p <0.01. For solutions for every set of 10,000 models, frequency of attaining E, M, or hybrid E/M state is plotted. Both plots (above, below) compare the frequency of attaining E, M, and hybrid E/M states in 10,000 control cases vs. 10,000 ZEB1-ESRP1 KD cases.

To further characterize the role of inhibition of ESRP1 by ZEB1 in mediating EMT/MET, we applied our recently developed computational method - Random Circuit Perturbation (RACIPE), [36] - to the miR-200/ZEB/ESRP1/CD44 circuit. For a given circuit topology, RACIPE generates an ensemble of mathematical models and identifies robust gene expression patterns and phenotypes that can be expected from that topology. Here, we used RACIPE to generate 10000 models for the circuit with an intact ZEB1-ESRP1 link (control case) and without this link (ZEB1-ESRP1 KD). Comparing the solutions of this ensemble of models from these both scenarios, attenuated inhibition of ESRP1 by ZEB1, i.e. effectively higher ESRP1 levels, led to decreased propensity to attain a mesenchymal phenotype and an increased propensity to adopt a hybrid E/M state, suggesting that ESRP1 overexpression may halt or reverse EMT (Figure 2C, S1).

Next, to experimentally test this prediction from our mathematical model, we treated MCF10A cells with TGFβ for 21 days, and then overexpressed ESRP1. Cells treated with TGFβ for 21 days underwent canonical morphological and biochemical changes associated with EMT such as spindle-shaped morphology and increased levels of ZEB1 (MCF10A-TGFβ-Ctrl) [28]. Upon overexpression of ESRP1 (MCF10A-TGFβ-ESRP1), these cells regained a cobblestone-shaped morphology and lost ZEB1 largely (Figure 3A, B). Thus, overexpression of EMT can revert EMT progression, an observation consistent with previous reports in using a tamoxifen-dependent activation of Twist-ER in mammary epithelial HMLE cells [30].

**Figure 3.**
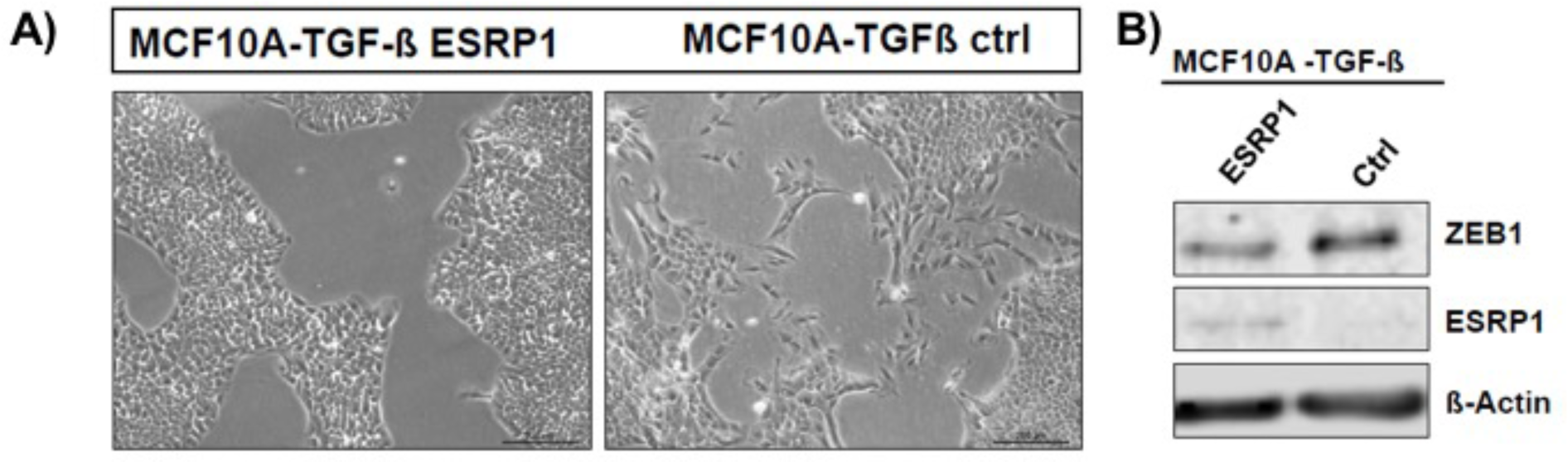
Overexpression of ESRP1 inhibits EMT in MCF10A cells. **A)** Bright-field microscopy images for MCF10A cells, transfected with a control vector (right) or an ESRP1 overexpression construct (left) - after treatment with TGFβ for 21 days. ***B)*** Western blot analysis showing changes in ESRP1 and ZEB1 levels in cells presented in A).

### ESRP1 knockdown destabilizes a hybrid epithelial/mesenchymal (E/M) phenotype

Next, we investigated how knockdown of ESRP1 can affect EMT. Our previous work has identified several ‘phenotypic stability factors’ (PSFs) such as OVOL2 and GRHL2 that can stabilize a hybrid E/M phenotype. They form a mutually inhibitory loop with ZEB1 [27], and their knockdown destabilizes a hybrid E/M phenotype both *in vitro* [27,37] and *in vivo* [38]. Because ESRP1 also forms an effective mutually inhibitory loop with ZEB1 by regulating the alternative splicing of CD44, we hypothesized that knockdown of ESRP1 may destabilize a hybrid E/M phenotype similarly.

To investigate this hypothesis, we knocked down ESRP1 via two independent siRNAs in H1975 - lung cancer cells that display a hybrid E/M phenotype stably over multiple passages *in vitro* under normal culturing conditions [27]. Untreated H1975 cells co-expressed E-cadherin (CDH1) and Vimentin (VIM) and showed nuclear localization of ZEB1 at a single-cell level (Figure 4, left 2 columns). Conversely, knockdown of ESRP1 substantially reduced E-cadherin and increased nuclear levels of ZEB1 (Figure 4A, right 2 columns), thus indicative of a transition towards a more mesenchymal phenotype. These observations go along with decrease of E-cadherin and increase of Vimentin and ZEB1 on protein and RNA levels (Figure 4B, S2) and an increased spindle-shape morphology (Figure 4C, S2) in ESRP1 knockdown cells, further supporting a shift towards a mesenchymal state.

**Figure 4.**
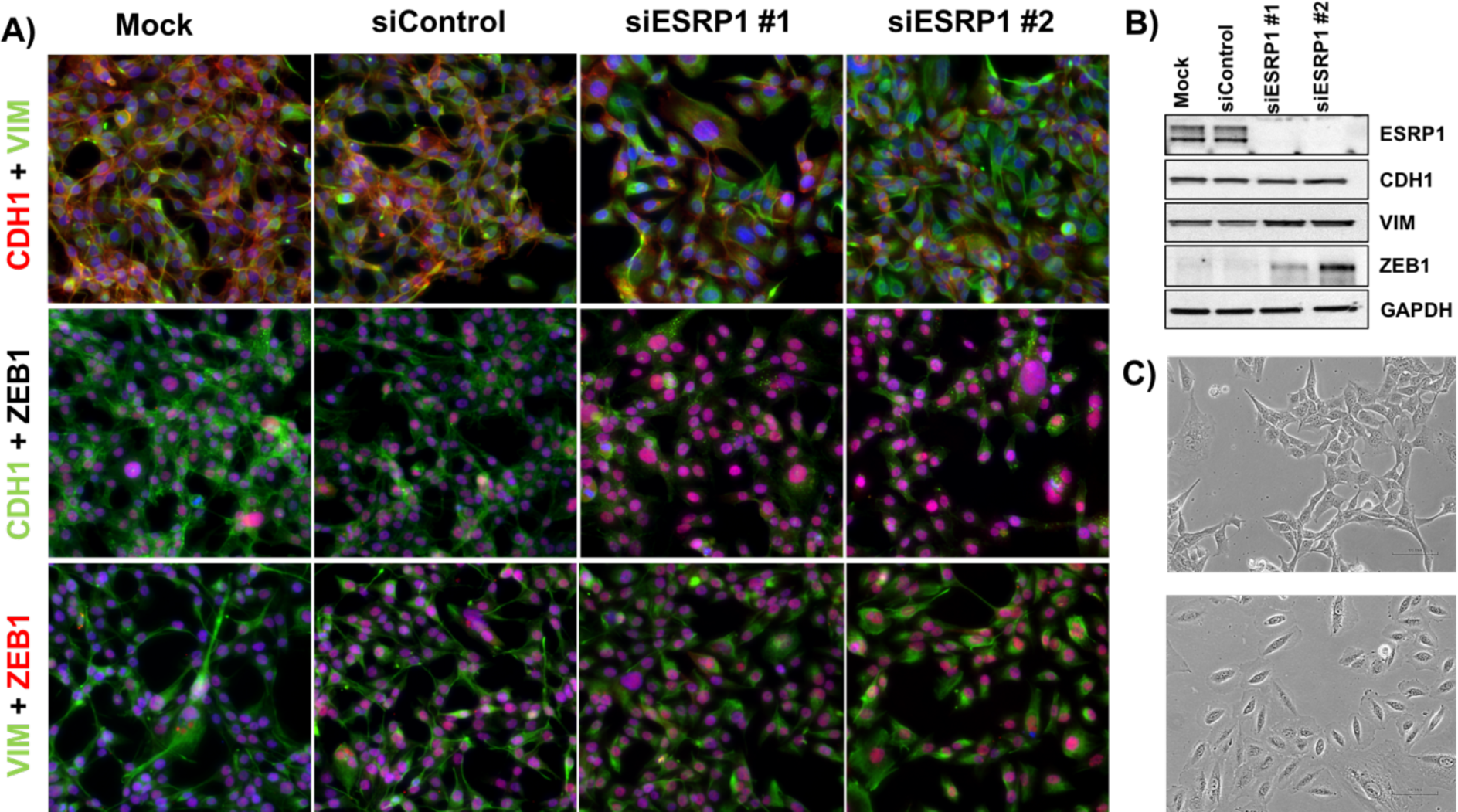
Knockdown of ESRP1 induces EMT in H1975 cells. **A)** Immunofluorescence images for hybrid E/M H1975 cells - mock (leftmost column), treated with control siRNA (2^nd^ column from the left), and treated with two independent siRNAs targeting ESRP1 (3^rd^, 4^th^ columns from the left). ***B)*** Western blot showing changes in ESRP1, CDH1, VIM, and ZEB1 levels. ***C)*** Bright-field microscopy images of mock (top) and si-ESRP1 #2 treated H1975 cells (bottom).

### ESRP1 expression correlates with prognosis of breast cancer patients

PSFs such as GRHL2 associate with poor prognosis in breast cancer [27,39], motivating us to investigate the correlation of ESRP1 with patient survival rates. Higher levels of ESRP1, and a higher ratio of ESRP1/HAS2 and ESRP1/ZEB1, correlate with poor relapse-free survival (RFS) and overall survival (OS) in multiple independent breast cancer datasets (Figure 5A-C, S3). These results suggest that the ZEB1/ESRP1/HAS2/CD44 axis may play important functional roles in breast cancer progression.

**Figure 5.**
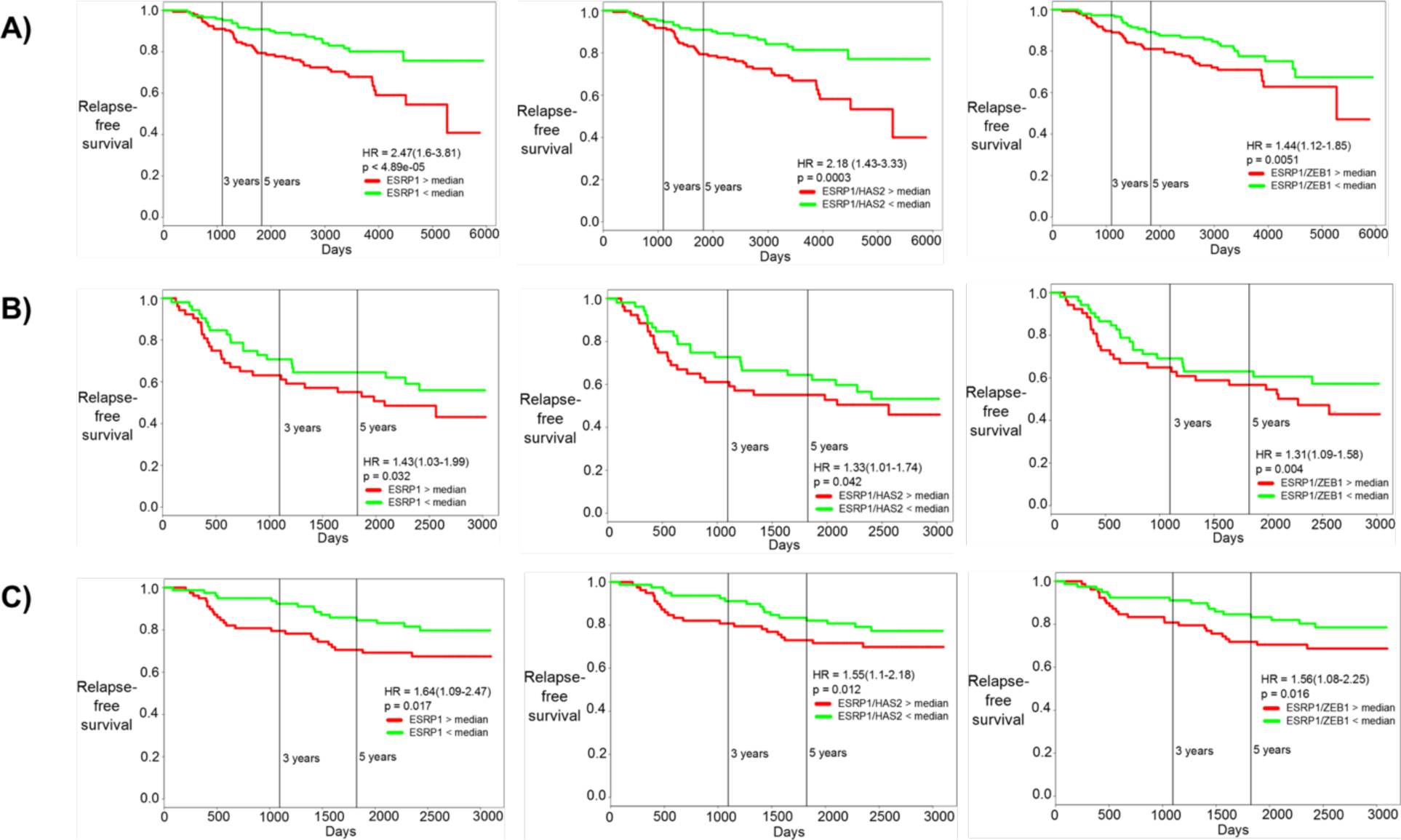
Correlation of ESRP1, ESRP1/HAS2, and ESRP1/ZEB1 with patient prognosis. Kaplan-Meier curves for relapse-free survival for ***A)*** GSE 17705, ***B)*** GSE 42568, and ***C)*** GSE 1456. HR stands for Hazard Ratio, i.e. the ratio of likelihood of relapse-free survival of patients with low ESRP1 as compared with those with high ESRP1. First column is for ESRP1, middle column is for ESRP1/HAS2 ratio, and right column is for ESRP1/ZEB1 ratio. Red curve shows patients with high ESRP1 levels, green curves patients with low ESRP1 levels (separated by median).

## Discussion

EMT is affected by multiple cross-wired cell-intrinsic and cell-extrinsic signals, including mechanical regulation through factors such as matrix stiffness [8,9,40]. Hyaluronan - also known as Hyaluronic Acid (HA) - is a proteoglycan that forms a scaffold for assembly of extracellular matrix (ECM). It mediates its metastasis-promoting effects largely by interacting with various isoforms of the cell surface receptor CD44 [41]. The relative abundance of these isoforms is regulated by an alternative splicing regulatory ESRP1 [29,30] that is altered during

EMT in multiple cancer types [26,30]. HA-CD44 interactions can activate ZEB1 - a key transcription factor driving EMT - which, in turn, not only inhibits ESRP1, but also upregulates HAS2 that synthesizes HA [28,29]. Put together, this set of feedback loops and complex interactions give rise to a highly nonlinear dynamic regulation of epithelial-hybrid-mesenchymal phenotypic transition that we characterized here through an integrated theoretical-experimental approach.

Our mathematical model predicts that the nonlinear dynamics emerging from these autocrine interconnected feedback loops can enable three phenotypes - epithelial (low ZEB1), mesenchymal (high ZEB1) and hybrid E/M (intermediate ZEB1). Multiple recent studies have identified a hybrid E/M phenotype at a single-cell level in cell lines, primary tumors, circulating tumor cells (CTCs) and metastatic tumors, across cancer types [42]. Cells in a hybrid E/M phenotype, in contrast to those in solely epithelial or mesenchymal states, can exhibit enhanced tumor-initiation potential, drug resistance and anoikis resistance *in vitro* and *in vivo* [43–49]. Moreover, many CTC clusters - the primary harbingers of metastasis [50] - exhibit a hybrid E/M signature [51]. These observations drive attention toward mechanisms maintaining a hybrid E/M phenotype [27,37,52–54].

Our experimental observations that ESRP1 overexpression can reverse EMT in MCF10A cells treated with TGFβ for 21 days, and that ESRP1 knockdown can drive EMT in H1975 cells propose ESRP1 as a potential factor that can stabilize a hybrid E/M phenotype. This idea is strengthened by our computational analysis using RACIPE that showing an enrichment of hybrid E/M phenotype when the inhibition of ESRP1 by ZEB1 is deleted (i.e. increased ESRP1 levels effectively). Finally, similar to the prognostic ability of GRHL2 and OVOL [27,39] - factors that can stabilize a hybrid E/M phenotype [25,27,37,38] - higher levels of ESRP1 have been associated with (a) poor OS in breast cancer, (b) poor 5-year progression-free survival (PFS) and OS in epithelial ovarian cancer samples [55,56], and (c) poor prognosis of distant metastasis in breast cancer samples [57]. Put together, these observations support a case for ESRP1 to be referred to as ‘phenotypic stability factor’ (PSF). However, including ESRP1 and its interconnections with the miR-200/ZEB circuit did not change the bifurcation diagram of the circuit, as was observed for other PSFs such as GRHL2, OVOL2, and ΔΝΡ63α [25,27,54].

Higher levels of ESRP1, as well as higher ratios of ESRP1/ZEB1 and/or ESRP1/HAS2 can predict poor survival in multiple breast cancer datasets. This observation indicates that both the ZEB1-ESRP1 and ZEB1-HAS2 axes may regulate ZEB1 levels simultaneously. Besides breast cancer, ZEB1 has been shown to directly repress ESRP1 in NSCLC [58,59], explaining the negative correlation between them in 22 NSCLC cell lines [60]. Our results showing that ESRP1 knockdown alters ZEB1 levels in H1975 - a hybrid E/M NSCLC cell line - suggest that this mutual inhibition between ESRP1 and ZEB1 may be functionally active in NSCLC. Increased ESRP1 can enrich for CD44v, thus consistent with the prognostic ability of ESRP1, higher levels of CD44v, but not high CD44s or total CD44 levels, are prognostic markers for distant metastases in lung and breast cancer [57].

CD44v can contribute to metastasis in multiple ways, e.g. by controlling the stability of cysteine transporter xCT that enables defense against enhanced oxidative stress [61]. Another example would be the production of an adhesive matrix - likely via anchoring to hyaluronan - to which tumor cells can attach during metastasis [62]. Contrarily, CD44s can also accelerate metastasis by increasing ZEB1 levels through HA-dependent and/or HA-independent pathways [28,29]. Thus, a fine-tuned balance of CD44s and CD44v isoforms may provide additional metastatic advantages. This balance is likely to correspond to a hybrid E/M phenotype which, as discussed above, may be more aggressive than cells in solely epithelial or mesenchymal states. Future studies would need sophisticated experimental models to understand spatiotemporal expression and function of CD44 isoforms during cancer progression.

In conclusion, our results demonstrate how two interconnected feedback loops of ZEB1 including ESRP1, HAS2, and CD44 govern the dynamics of EMT/MET and enable the existence of a stable hybrid E/M phenotype. ZEB1 can not only enforce its own expression through these loops, but also alter its microenvironment, thus enhancing non-cell autonomous effects of EMT. Similar non-cell autonomous effects can lead to a cooperative behavior between epithelial and mesenchymal cells in establishing metastases [44,63,64]. Moreover, these non-cell autonomous effects of EMT mediated by ZEB1 may help offer a plausible explanation why metastasis can be blunted largely by knockout of ZEB1 [65], but not necessarily by that of SNAIL or TWIST [66].

## Author contributions

Designed research: M.K.J., B-T.P.

Performed research: M.K.J., B-T.P., S.C.T., D.J.

Analyzed data: M.K.J., B-T.P., S.C.T., D.J., S.M., M.S., T.B., J.M., H. L.

Wrote manuscript: M.K.J., B-T.P., D.J., J.M., H.L.

Supervised research: J.M., H.L.

## Acknowledgements

This work is supported by National Science Foundation (NSF) Center for Theoretical Biological Physics (NSF PHY-1427654) and NSF DMS-1361411. M.K.J. is also supported by a training fellowship from the Gulf Coast Consortia on Computational Cancer Biology Training Program (CPRIT Grant No. RP170593), and would like to thank Cindy Farach-Carson for useful discussions. J.M. is supported by a grant from the Interdisciplinary Center for Clinical Research within the faculty of Medicine at the RWTH Aachen University (AP 1-4) and by a Eurostars grant (E11497).

**Abbreviations**
CCLE: Cancer Cell Line Encyclopedia
CTC: Circulating Tumor Cell
ECM: extracellular matrix
EMT: Epithelial Mesenchymal Transition
ESRP1: Epithelial Splicing Regulatory Protein 1
HA: Hyaluronic Acid
HAS2: Hyaluronan Acid Synthase 2
MET: Mesenchymal Epithelial Transition
NSCLC: non-small cell lung cancer
RACIPE: Random Circuit Perturbation

## Materials and Methods

### Cell Culture

Cell line MCF10A was purchased from American Type Culture Collection (ATCC). Cells were cultured in DMEM/F12 (Invitrogen, Karlsruhe, Germany, 31331) containing 5% horse serum (Life Technologies, Darmstadt, Germany, 16050122), 20 ng/ml EGF (R&D Systems, Wiesbaden, Germany, 236EG200), 0.5 mg/ml hydrocortisone (Sigma, Taufkirchen, Germany, H0888#1G), 0.1 mg/ml cholera toxin (Sigma, Taufkirchen, Germany, C-8052) and 10 mg/ml insulin (Invitrogen, Karlsruhe, Germany, 12585-014). Induction of EMT by TGFb1 (PeproTech, Hamburg, Germany, 100-21) in MCF10A cells was performed by adding daily 5 ng/ml TGFb1 to the medium of MCF10A cultures for 21 days. Medium was replaced every second day. After 21 days, MCF10A ESRP1 cells were generated by lentiviral infection with an ESRP1 overexpression and control construct (pCDH-CMV-(ESRP1)-MCS-EF1-copGFP).

H1975 cells were cultured in RPMI 1640 medium containing 10% fetal bovine serum and 1% penicillin/streptomycin cocktail (Thermo Fisher Scientific, Waltham, MA). Cells were transfected at a final concentration of 50 nM siRNA using Lipofectamine RNAiMAX (Thermo Fisher Scientific) according to the manufacturer’s instructions using following siRNAs: siControl (Silencer Select Negative Control No. 1, Thermo Fisher Scientific), siESRP1 #1 (SASI_Hs01_00062865, Sigma-Aldrich, St.Louis, MO), siESRP1 #2 (SASI_Hs02_00308155, Sigma-Aldrich, St.Louis, MO),).

### Western blotting analysis and immunoflourescence

MCF10A cCells were rinsed once in PBS and lysed in TLB. 30 μg of protein was separated by SDS-PAGE (10%) for 1 h, 150 V and transferred to a nitrocellulose membrane by wet blotting in transfer buffer for 2 h, 300 mA at 4°C. Membranes were immersed in Antigen pretreatment solution (SuperSignal Westernblot Enhancer, Thermo Scientific) for 10 min and blocked in 5% skim milk/ TBST) for 30 min at room temperature. Primary antibody incubation was carried out in Primary antibody Diluent (SuperSignal Westernblot Enhancer, Thermo, 46641) overnight at 4°C. After washing in TBST, membrane was incubated with HRP-conjugated secondary antibody in 5% skim milk/TBST for 1 h at RT. Detection was carried out using SuperSignal West Femto Maximum Sensitivity Substrate (Thermo, 34094) or ECL Prime Westenblot Detection Reagent (Amersham, RPN2232) and a ChemiDoc imaging system (BioRad). Quantification was performed where appropriate using ImageJ and presented normalized to β-Actin levels.

H1975 Cells were lysed in RIPA lysis assay buffer (Pierce) supplemented with protease and phosphatase inhibitor. The samples were separated on a 4-15% SDS-polyacrylamide gel (Biorad). After transfer to PVDF membrane, probing was carried out with primary antibodies and subsequent secondary antibodies. Primary antibodies were purchased from the following commercial sources: anti-CDH1 (1:1000; Cell Signaling Technology), anti-vimentin (1:1000; Cell Signaling Technology), anti-ESRP1 (0.4ug/ml; Sigma), anti-ZEB1 (1:200; Abcam) and anti-GAPDH (1:10,000; Abcam). Membranes were exposed using the ECL method (GE Healthcare) according to the manufacturer’s instructions. For immunofluorescence, cells were fixed in 4% paraformaldehyde, permeabilized in 0.2% Triton X-100, and then stained with anti-CDH1 (1:100; Abcam), anti-vimentin (1:100; Cell Signaling Technology) and anti-ZEB1 (1:200; Abcam). The primary antibodies were then detected with Alexa congugated secondary antibodies (Life technologies). Nuclei were visualized by co-staining with DAPI.

### Transfection of plasmid DNA

Plasmid DNA transfection was done by using the FugeneHD transfection reagent (Promega, E2311) according to the manufacturer’s instructions and harvested 72 h afterwards for protein analysis.

### Antibodies

The following antibodies and dilutions were used for Western blotting: mouse anti-β -Actin (Sigma, A5441; 1:5,000), rabbit anti-ZEB1 (Sigma, HPA027524; 1:5,000), as well as HRP-coupled goat anti-rabbit IgG (Dianova, 111-035-003; 1:25,0000) and goat anti-mouse IgG (Dianova, 115-035-003; 1:25,000), mouse anti-ESRP1 (Abnova, Heidelberg, Germany, ab140671; 1:500)

### RT-PCR

Total RNA was isolated following manufacturer’s instructions using RNAeasy kit (Qiagen). cDNA was prepared using iScript gDNA clear cDNA synthesis kit (Bio-Rad). A TaqMan PCR assay was performed with a 7500 Fast Real-Time PCR System using TaqMan PCR master mix, commercially available primers, and FAM™-labeled probes for CDH1, VIM, ZEB1, ESRP1 and VIC™-labeled probes for 18S, according to the manufacturer’s instructions (Life Technologies). Each sample was run in triplicate. Ct values for each gene were calculated and normalized to Ct values for 18S (ΔCt). The ΔΔCt values were then calculated by normalization to the ΔCt value for control.

### Mathematical model

Section S1 contains all details of mathematical model development, analysis, and parameter estimates.

## References

1. Nieto MA (2013) Epithelial plasticity: a common theme in embryonic and cancer cells. Science 342: 1234850.

2. Tsai JH, Yang J (2013) Epithelial-mesenchymal plasticity in carcinoma metastasis. Genes Dev 27: 2192–2206.

3. Jolly MK, Ware KE, Gilja S, Somarelli JA, Levine H (2017) EMT and MET: necessary or permissive for metastasis? Mol Oncol 11: 755–769.

4. Singh A, Settleman J (2011) EMT, cancer stem cells and drug resistance: an emerging axis of evil in the war on cancer. Oncogene 29: 4741–4751.

5. Tripathi SC, Peters HL, Taguchi A, Katayama H, Wang H, Momin A, Jolly MK, Celiktas M, Rodriguez-Canales J, Liu H, et al. (2016) Immunoproteasome deficiency is a feature of non-small cell lung cancer with a mesenchymal phenotype and is associated with a poor outcome. Proc Natl Acad Sci 113: E1555–64.

6. Wellner U, Schubert J, Burk UC, Schmalhofer O, Zhu F, Sonntag A, Waldvogel B, Vannier C, Darling D, zur Hausen A, et al. (2009) The EMT-activator ZEB1 promotes tumorigenicity by repressing stemness-inhibiting microRNAs. Nat Cell Biol 11: 1487–1495.

7. Jolly MK, Huang B, Lu M, Mani SA, Levine H, Ben-Jacob E (2014) Towards elucidating the connection between epithelial - mesenchymal transitions and stemness. J R Soc Interface 11: 20140962.

8. Rice AJ, Cortes E, Lachowski D, Cheung BCH, Karim SA, Morton JP, Hernández AR (2017) Matrix stiffness induces epithelial - mesenchymal transition and promotes chemoresistance in pancreatic cancer cells. Oncogenesis 6: e352.

9. Wei SC, Fattet L, Tsai JH, Guo Y, Pai VH, Majeski HE, Chen AC, Sah RL, Taylor SS, Engler AJ, et al. (2015) Matrix stiffness drives epithelial-mesenchymal transition and tumour metastasis through a TWIST1-G3BP2 mechanotransduction pathway. Nat Cell Biol 17: 678–688.

10. Zhang L, Huang G, Li X, Zhang Y, Jiang Y, Shen J, Liu J, Wang Q, Zhu J, Feng X, et al. (2013) Hypoxia induces epithelial-mesenchymal transition via activation of SNAI1 by hypoxia-inducible factor -1α in hepatocellular carcinoma. BMC Cancer 13: 108.

11. De Craene B, Berx G (2013) Regulatory networks defining EMT during cancer initiation and progression. Nat Rev Cancer 13: 97–110.

12. Liu Y, Lu X, Huang L, Wang W, Jiang G, Dean KC, Clem B, Telang S, Jenson AB, Cuatrecasas M, et al. (2014) Different thresholds of ZEB1 are required for Ras-mediated tumour initiation and metastasis. Nat Commun 5: 5660.

13. Meidhof S, Brabletz S, Lehmann W, Preca B-T, Mock K, Ruh M, Schüler J, Berthold M, Weber A, Burk U, et al. (2015) ZEB1-associated drug resistance in cancer cells is reversed by the class I HdAC inhibitor mocetinostat. EMBO Mol Med 7: 831–847.

14. Bronsert P, Kohler I, Timme S, Kiefer S, Werner M, Schilling O, Vashist Y, Makowiec F, Brabletz T, Hopt UT, et al. (2014) Prognostic significance of Zinc finger E-box binding homeobox 1 (ZEB1) expression in cancer cells and cancer-associated fibroblasts in pancreatic head cancer. Surgery 156: 97–108.

15. Somarelli JA, Shelter S, Jolly MK, Wang X, Bartholf Dewitt S, Hish AJ, Gilja S, Eward WC, Ware KE, Levine H, et al. (2016) Mesenchymal-epithelial transition in sarcomas is controlled by the combinatorial expression of miR-200s and GRHL2. Mol Cell Biol 36: 2503–2513.

16. Sundararajan V, Gengenbacher N, Stemmler MP, Kleemann JA, Brabletz T BS (2015) The ZEB1/miR-200c feedback loop regulates invasion via actin interacting proteins MYLK and TKS5. Oncotarget 6: 27083–27096.

17. Burk U, Schubert J, Wellner U, Schmalhofer O, Vincan E, Spaderna S, Brabletz T (2008) A reciprocal repression between ZEB1 and members of the miR-200 family promotes EMT and invasion in cancer cells. EMBO Rep 9: 582–589.

18. Bracken CP, Gregory PA, Kolesnikoff N, Bert AG, Wang J, Shannon MF, Goodall GJ (2008) A double-negative feedback loop between ZEB1-SIP1 and the microRNA-200 family regulates epithelial-mesenchymal transition. Cancer Res 68: 7846–7854.

19. Brabletz S, Brabletz T (2010) The ZEB/miR-200 feedback loop–a motor of cellular plasticity in development and cancer? EMBO Rep 11: 670–677.

20. Gregory PA, Bracken CP, Smith E, Bert AG, Wright J a, Roslan S, Morris M, Wyatt L, Farshid G, Lim Y-Y, et al. (2011) An autocrine TGF-beta/ZEB/miR-200 signaling network regulates establishment and maintenance of epithelial-mesenchymal transition. Mol Biol Cell 22: 1686–1698.

21. Tian X-J, Zhang H, Xing J (2013) Coupled Reversible and Irreversible Bistable Switches Underlying TGFβ-induced Epithelial to Mesenchymal Transition. Biophys J 105: 1079–1089.

22. Zhang J, Tian X-J, Zhang H, Teng Y, Li R, Bai F, Elankumaran S, Xing J (2014) TGF-β-induced epithelial-to-mesenchymal transition proceeds through stepwise activation of multiple feedback loops. Sci Signal 7: ra91.

23. Lu M, Jolly MK, Levine H, Onuchic JN, Ben-Jacob E (2013) MicroRNA-based regulation of epithelial-hybrid-mesenchymal fate determination. Proc Natl Acad Sci 110: 18144–18149.

24. Li C, Hong T, Nie Q (2016) Quantifying the landscape and kinetic paths for epithelial-mesenchymal transition from a core circuit. Phys Chem Chem Phys 18: 17949–17956.

25. Jia D, Jolly MK, Boareto M, Parsana P, Mooney SM, Pienta KJ, Levine H, Ben-Jacob E (2015) OVOL guides the epithelial-hybrid-mesenchymal transition. Oncotarget 6: 15436–15448.

26. Jia D, Jolly MK, Tripathi SC, Hollander P Den, Huang B, Lu M, Celiktas M, Ramirez-Pena E, Ben-Jacob E, Onuchic JN, et al. (2017) Distinguishing Mechanisms Underlying EMT Tristability. Cancer Converg 1: 2.

27. Jolly MK, Tripathi SC, Jia D, Mooney SM, Celiktas M, Hanash SM, Mani SA, Pienta KJ, Ben-Jacob E, Levine H (2016) Stability of the hybrid epithelial/mesenchymal phentoype. Oncotarget 7: 27067–27084.

28. Preca B-T, Bajdak K, Mock K, Lehmann W, Sundararajan V, Bronsert P, Matzge-Ogi A, Orian-Rousseau V, Brabletz S, Brabletz T, et al. (2017) A novel ZEB1/HAS2 positive feedback loop promotes EMT in breast cancer. Oncotarget 8: 11530–11543.

29. Preca B-T, Bajdak K, Mock K, Sundararajan V, Pfannstiel J, Maurer J, Wellner U, Hopt UT, Brummer T, Brabletz S, et al. (2015) A self-reinforcing CD44s/ZEB1 feedback loop maintains EMT and stemness properties in cancer cells. Int J Cancer 137: 2566–2577.

30. Brown RL, Reinke LM, Damerow MS, Perez D, Chodosh L a., Yang J, Cheng C (2011) CD44 splice isoform switching in human and mouse epithelium is essential for epithelial-mesenchymal transition and breast cancer progression. J Clin Invest 121: 1064–1074.

31. Cieply B, Koontz C, Frisch SM (2015) CD44S-hyaluronan interactions protect cells resulting from EMT against anoikis. Matrix Biol 48: 55–65.

32. Paoli P, Giannoni E, Chiarugi P (2013) Anoikis molecular pathways and its role in cancer progression. Biochim Biophys Acta - Mol Cell Res 1833: 3481–3498.

33. Liu S, Cheng C (2017) Akt signaling is sustained by a CD44 splice isoform-mediated positive feedback loop. Cancer Res 77: 3791–3801.

34. Huang S, Guo YP, May G, Enver T (2007) Bifurcation dynamics in lineage-commitment in bipotent progenitor cells. Dev Biol 305: 695–713.

35. Jia D, Jolly MK, Harrison W, Boareto M, Ben-Jacob E, Levine H (2017) Operating principles of tristable circuits regulating cellular differentiation. Phys Biol 4: 35007.

36. Huang B, Lu M, Jia D, Ben-Jacob E, Levine H, Onuchic JN (2017) Interrogating the topological robustness of gene regulatory circuits by randomization. PLoS Comput Biol 13: e1005456.

37. Hong T, Watanabe K, Ta CH, Villarreal-Ponce A, Nie Q, Dai X (2015) An Ovol2-Zeb1 Mutual Inhibitory Circuit Governs Bidirectional and Multi-step Transition between Epithelial and Mesenchymal States. PLOS Comput Biol 11: e1004569.

38. Watanabe K, Villarreal-Ponce A, Sun P, Salmans ML, Fallahi M, Andersen B, Dai X (2013) Mammary morphogenesis and regeneration require the inhibition of EMT at terminal end buds by Ovol2 transcriptional repressor. Dev Cell 29: 59–74.

39. Mooney SM, Talebian V, Jolly MK, Jia D, Gromala M, Levine H, McConkey BJ (2017) The GRHL2/ZEB Feedback Loop - A Key Axis in the Regulation of EMT in Breast Cancer. J Cell Biochem in press.

40. Kumar S, Das A, Sen S (2014) Extracellular matrix density promotes EMT by weakening cell-cell adhesions. Mol Biosyst 10: 838–850.

41. Bourguignon LYW, Earle C, Shiina M (2017) Activation of Matrix Hyaluronan-Mediated CD44 Signaling, Epigenetic Regulation and Chemoresistance in Head and Neck Cancer Stem Cells. Int J Mol Sci 18: 1849.

42. Jolly MK, Boareto M, Huang B, Jia D, Lu M, Ben-Jacob E, Onuchic JN, Levine H (2015) Implications of the hybrid epithelial/mesenchymal phenotype in metastasis. Front Oncol 5: 155.

43. Biddle A, Gammon L, Liang X, Costea DE, Mackenzie IC (2016) Phenotypic Plasticity Determines Cancer Stem Cell Therapeutic Resistance in Oral Squamous Cell Carcinoma. EBioMedicine 4: 138–145.

44. Grosse-Wilde A, Fouquier d’ Herouei A, McIntosh E, Ertaylan G, Skupin A, Kuestner RE, del Sol A, Walters K-A, Huang S (2015) Stemness of the hybrid epithelial/mesenchymal state in breast cancer and its association with poor survival. PLoS One 10: e0126522.

45. Goldman A, Majumder B, Dhawan A, Ravi S, Goldman D, Kohandel M, Majumder PK, Sengupta S (2015) Temporally sequenced anticancer drugs overcome adaptive resistance by targeting a vulnerable chemotherapy-induced phenotypic transition. Nat Commun 6: 6139.

46. Huang RY-J, Wong MK, Tan TZ, Kuay KT, Ng a HC, Chung VY, Chu Y-S, Matsumura N, Lai H-C, Lee YF, et al. (2013) An EMT spectrum defines an anoikis-resistant and spheroidogenic intermediate mesenchymal state that is sensitive to e-cadherin restoration by a src-kinase inhibitor, saracatinib (AZD0530). Cell Death Dis 4: e915.

47. Andriani F, Bertolini G, Facchinetti F, Baldoli E, Moro M, Casalini P, Caserini R, Milione M, Leone G, Pelosi G, et al. (2016) Conversion to stem-cell state in response to microenvironmental cues is regulated by balance between epithelial and mesenchymal features in lung cancer cells. Mol Oncol 10: 253–271.

48. Jolly MK, Jia D, Boareto M, Mani SA, Pienta KJ, Ben-Jacob E, Levine H (2015) Coupling the modules of EMT and stemness: A tunable ‘stemness window’ model. Oncotarget 6: 25161–25174.

49. Bierie B, Pierce SE, Kroeger C, Stover DG, Pattabiraman DR, Thiru P, Liu Donaher J, Reinhardt F, Chaffer CL, Keckesova Z, et al. (2017) Integrin-β4 identifies cancer stem cell-enriched populations of partially mesenchymal carcinoma cells. Proc Natl Acad Sci 114: E2337–2346.

50. Cheung KJ, Ewald AJ (2016) A collective route to metastasis: Seeding by tumor cell clusters. Science 352: 167–169.

51. Sarioglu AF, Aceto N, Kojic N, Donaldson MC, Zeinali M, Hamza B, Engstrom A, Zhu H, Sundaresan TK, Miyamoto DT, et al. (2015) A microfluidic device for label-free, physical capture of circulating tumor cell clusters. Nat Methods 12: 685–691.

52. Bocci F, Jolly MK, Tripathi SC, Aguilar M, Onuchic N, Hanash SM, Levine H, Levine H (2017) Numb prevents a complete epithelial - mesenchymal transition by modulating Notch signalling. J R Soc Interface 14: 20170512.

53. Dang TT, Esparza MA, Maine EA, Westcott JM, Pearson GW (2015) ΔNp63α promotes breast cancer cell motility through the selective activation of components of the Epithelial-to-Mesenchymal Transition program. Cancer Res 75: 3925–3935.

54. Jolly MK, Boareto M, Debeb BG, Aceto N, Farach-Carson MC, Woodward WA, Levine H (2017) Inflammatory Breast Cancer: a model for investigating cluster-based dissemination. NPJ Breast Cancer 3: 21.

55. Jeong HM, Han J, Lee SH, Park H, Lee HJ, Choi J, Lee YM, Choi Y, Shin YK, Kwon MJ (2017) ESRP1 is overexpressed in ovarian cancer and promotes switching from mesenchymal to epithelial phenotype in ovarian cancer cells. Oncogenesis 6: e389.

56. Chen L, Yao Y, Sun L, Zhou J, Miao M, Luo S, Deng G, Li J, Wang J, Tang J (2017) Snail Driving Alternative Splicing of CD44 by ESRP1 Enhances Invasion and Migration in Epithelial Ovarian Cancer. Cell Physiol Biochem 43: 2489–2504.

57. Hu J, Li G, Zhang P, Zhuang X, Hu G (2017) A CD44v + subpopulation of breast cancer stem-like cells with enhanced lung metastasis capacity. Cell Death Dis 8: e2679.

58. Larsen JE, Nathan V, Osborne JK, Farrow RK, Deb D, Sullivan JP, Dospoy PD, Augustyn A, Hight SK, Sato M, et al. (2016) ZEB1 drives epithelial-to-mesenchymal transition in lung cancer. J Clin Invest 126: 3219–3235.

59. Roche J, Nasarre P, Gemmill R, Baldys A, Pontis J, Ait-si-ali S, Drabkin H (2013) Global Decrease of Histone H3K27 Acetylation in ZEB1-Induced Epithelial to Mesenchymal Transition in Lung Cancer Cells. Cancers (Basel) 5: 334–356.

60. Gemmill RM, Roche J, Potiron VA, Nasarre P, Mitas M, Coldren CD, Helfrich B, Garrett-Mayer E, Bunn PA, Drabkin HA (2011) ZEB1-responsive genes in non-small cell lung cancer. Cancer Lett 300: 66–78.

61. Yae T, Tsuchihashi K, Ishimoto T, Motohara T, Yoshikawa M, Yoshida GJ, Wada T, Masuko T, Mogushi K, Tanaka H, et al. (2012) Alternative splicing of CD44 mRNA by ESRP1 enhances lung colonization of metastatic cancer cell. Nat Commun 3: 883.

62. Klingbeil P, Marhaba R, Jung T, Kirmse R, Ludwig T, Zoller M (2009) CD44 Variant Isoforms Promote Metastasis Formation by a Tumor Cell-Matrix Cross-talk That Supports Adhesion and Apoptosis Resistance. Mol Cancer Res 7: 168–180.

63. Tsuji T, Ibaragi S, Shima K, Hu MG, Katsurano M, Sasaki A, Hu GF (2008) Epithelial-mesenchymal transition induced by growth suppressor p12 CDK2-AP1 promotes tumor cell local invasion but suppresses distant colony growth. Cancer Res 68: 10377–10386.

64. Neelakantan D, Zhou H, Oliphant MUJ, Zhang X, Simon LM, Henke DM, Shaw CA, Wu MF, Hilsenbeck SG, White LD, et al. (2017) EMT cells increase breast cancer metastasis via paracrine GLI activation in neighbouring tumour cells. Nat Commun 8: 15773.

65. Krebs AM, Mitschke J, Losada ML, Schmalhofer O, Boerries M, Busch H, Boettcher M, Mougiakakos D, Reichardt W, Bronsert P, et al. (2017) The EMT-activator Zeb1 is a key factor for cell plasticity and promotes metastasis in pancreatic cancer. Nat Cell Biol 19: 518–529.

66. Zheng X, Carstens JL, Kim J, Scheible M, Kaye J, Sugimoto H, Wu C-C, LeBleu VS, Kalluri R (2015) Epithelial-to-mesenchymal transition is dispensable for metastasis but induces chemoresistance in pancreatic cancer. Nature 527: 525–530.

